# See-Star: a versatile hydrogel-based protocol for clearing large, opaque and calcified marine invertebrates

**DOI:** 10.1101/2024.03.16.585344

**Authors:** D. N. Clarke, L. Formery, C. J. Lowe

## Abstract

Studies of morphology and developmental patterning in adult stages of many invertebrates are hindered by opaque structures, such as shells, skeletal elements, and pigment granules that block or refract light and necessitate sectioning for observation of internal features. An inherent challenge in studies relying on surgical approaches is that cutting tissue is semi-destructive, and delicate structures, such as axonal processes within neural networks, are computationally challenging to reconstruct once disrupted. To address this problem, we developed See-Star, a hydrogel-based tissue clearing protocol to render the bodies of opaque and calcified invertebrates optically transparent while preserving their anatomy in an unperturbed state, facilitating molecular labeling and observation of intact organ systems. The resulting protocol can clear large (>1 cm^3^) specimens to enable deep-tissue imaging, and is compatible with molecular techniques, such as immunohistochemistry and *in situ* hybridization to visualize protein and mRNA localization. To test the utility of this method, we performed a whole mount imaging study of intact nervous systems in juvenile echinoderms and molluscs, and demonstrate that See-Star allows for comparative studies to be extended far into development facilitating insights into the anatomy of juveniles and adults that are usually not amenable to whole mount imaging.

## Introduction

There is a wealth of anatomical diversity across metazoans that can inform questions on the evolution of animal body plans and organ systems. However, modern research in developmental biology has largely focused on a subset of model organisms, introducing some level of bias in our understanding of animal macroevolution. A way to solve this issue is through comparative observations in phylogenetically diverse taxa to expand our understanding of the evolution of organs systems and body plans (Brenowitz & Zakon, 2015; Hejnol & Lowe, 2015). Development of broadly applicable, adaptable research approaches is crucial for achieving this goal.

A pivotal challenge in biological research is visualizing cells or molecules within their native tissue context, as well as imaging entire tissues and organisms. This often requires dissection or serial sectioning of tissues followed by reconstruction, which is labor-intensive. Such tasks are particularly demanding for complex structures like the nervous system, where it is exceptionally hard to piece together fragmented neurons and intricate neural projections. While sophisticated microscopy techniques can provide high-resolution images of thin samples, achieving detailed images of entire nervous systems, or other organ systems, across whole organisms continues to be a significant obstacle, especially outside of well-characterized model organisms. Compounding this difficulty is the scattering of light due to varying refractive indices (RIs) among biological molecules—such as water, lipids, and proteins—and the absorption of light by endogenous pigments and calcified structures (Tainaka et al., 2016). For some organisms, such as shelled or heavily calcified marine invertebrates, this challenge is insurmountable, making the observation of internal structures in adult stages impossible without dissection or sectioning. In the context of evolutionary developmental biology, this has caused studies to often be restricted to developmental stages that are optically clear, such as embryos and larvae, due to imaging limitations.

Recent advances in tissue clearing methodologies have largely resolved the challenge of varying RIs to enable three-dimension observation of intact mammalian tissues in unprecedented detail, but these tools have received limited use and adaptation in non-mammalian systems (Azaripour et al., 2016; Tainaka et al., 2016). These methods typically aim to: (1) remove lipids from optically dense, fatty tissues like the central nervous system, and (2) match the RI within tissues through the use of specialized media to minimize light scattering. Solvent-based strategies, such as 3DISCO (Ertürk et al., 2012) and EZ Clear (Hsu et al., 2022), and strategies relying on detergents or other hydrophilic reagents, such as CUBIC (Susaki et al., 2014, 2015), CLARITY (Chung et al., 2013), and SeeDB (Ke et al., 2013), for delipidation have enabled molecular interrogation of intact neural circuits within mammalian brains (reviewed in Tainaka et al., 2016). Hydrogel-based strategies, like CLARITY, in which tissues are crosslinked to a polyacrylamide matrix through fixation and gelation, can provide a structural scaffold to support delicate tissues, and have been adapted for tissue clearing of decalcified bones (Greenbaum et al., 2017). While some efforts have been made to adapt tissue clearing protocols to non-model systems (Konno & Okazaki, 2018; Pende et al., 2020), none have demonstrated the capacity to preserve morphology in delicate, heavily calcified organisms.

Here, we describe a novel, robust, and simple method for visualizing organ systems in calcified and pigmented marine invertebrates that combines hydrogel crosslinking, decalcification, and tissue clearing. This method, which we term See-Star, preserves tissue integrity of delicate, highly calcified marine invertebrates following chemical decalcification. We demonstrate the efficacy of the See-Star protocol on two phyla of highly pigmented and calcified animals, echinoderms and molluscs. We show that See-Star is compatible with common molecular techniques, including immunohistochemistry (IHC) and *in situ* hybridization (ISH). See-Star enables imaging of molecular labels across scales in large samples, from high-resolution imaging of cellular structures up to whole-body imaging of organ systems. Thus, See-Star is a versatile approach that will enable molecular characterization of intact organ systems in developmental stages that are usually challenging for whole mount imaging, such as juveniles or young adults, in a variety of non-model organisms.

## Results

### See-Star renders calcified invertebrates transparent while preserving tissue integrity

Heavily calcified marine invertebrates often become very fragile after decalcification. We reasoned that a hydrogel-crosslinking fixation protocol could provide adequate support to maintain tissue structure through this process. Our goal was to create a protocol that combines robust fixation to preserve delicate tissues with decalcification to eliminate opaque skeletal elements, while also achieving efficient tissue clearing. We hypothesized that using distinct buffers for fixation and gelation would offer the flexibility to independently optimize each step (Fig. 1A). Unlike most hydrogel-based clearing methods, which use a single buffer for both the fixation and gelation steps, our approach allows for this necessary flexibility.

**Figure 1:**
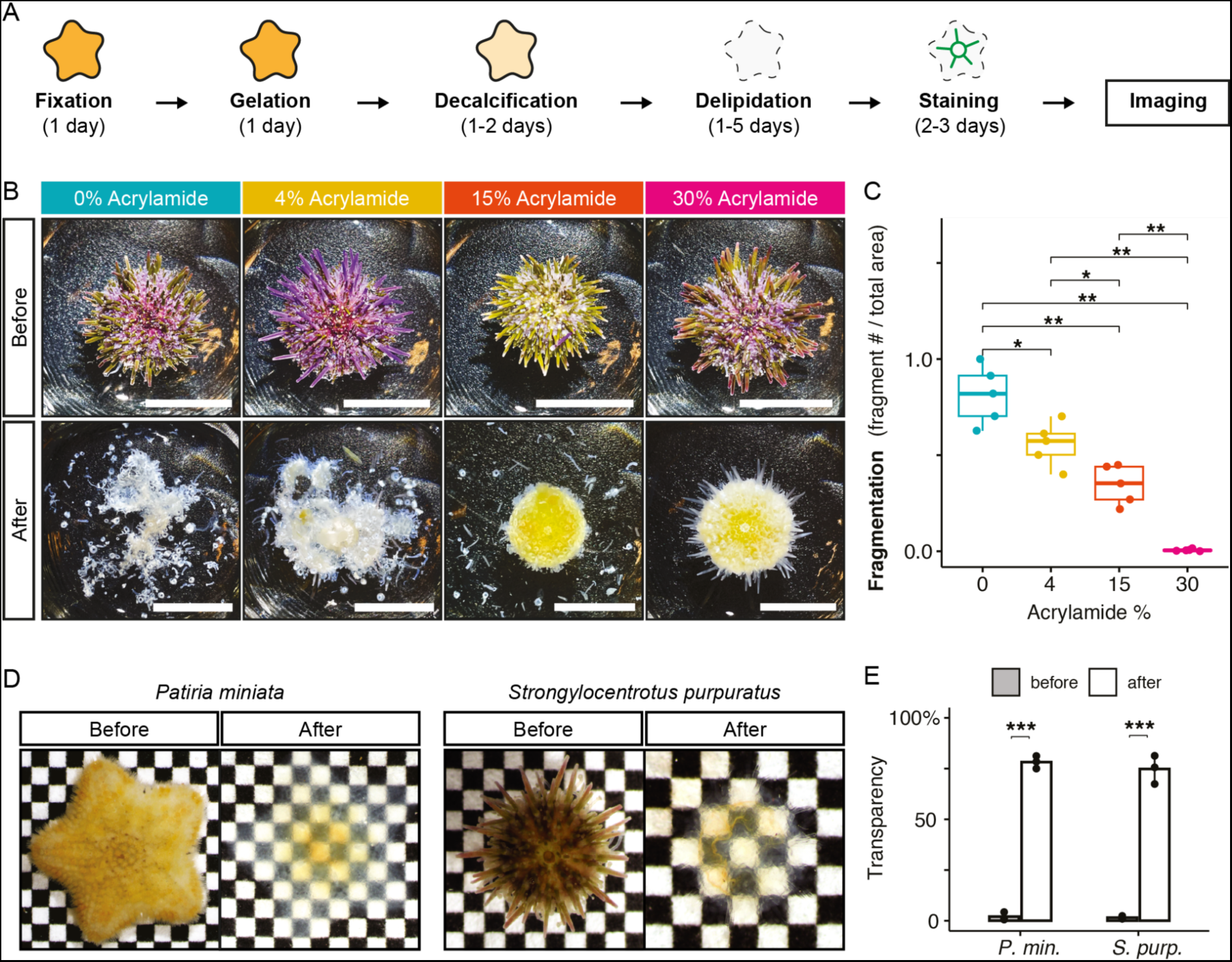
See-Star renders calcified invertebrates transparent while preserving tissue integrity A, Principle of See-Star clearing procedure, with approximate timing given for each step. **B,** Representative images of *Strongylocentrotus purpuratus* juveniles before and after preparation with varying concentrations of acrylamide. Scale bar = 1 cm. **C,** Quantification of sample fragmentation observed across varying concentrations of acrylamide. n = 5 per condition. **D,** Brightfield images of representative cleared echinoderm juveniles (left: *Patiria miniata*, asteroid; right: *S. purpuratus*, echinoid) imaged before clearing (following fixation) and after clearing (before imaging). Squares = 600 µm. **E,** Quantification of transparency before and after the clearing process for the species shown in **D**. n = 3 per condition. In **C**, **E**, statistical significance determined by Wilcoxon test; p < 0.05 = *, p < 0.01 = **, p < 0.001 = ***.

In preliminary experiments, we observed that the standard 4% acrylamide fixation used in CLARITY, and related hydrogel-based clearing protocols, was insufficient to preserve tissue integrity in adult echinoderms following decalcification (Fig. 1B). We hypothesized that increasing the acrylamide concentration in the fixation step would increase tissue robustness. To test this, we prepared samples of juvenile purple sea urchins, *Stronglyocentrotus purpuratus*, across a range of acrylamide concentrations. We found that acrylamide concentration correlated with tissue integrity: samples prepared with no or low percentages of acrylamide were severely fragmented following decalcification, whereas samples with the highest percentages of acrylamide survived decalcification and delipidation intact (Fig. 1B, C). To verify that samples fixed with 30% acrylamide could still be rendered optically transparent, we next measured the transparency of samples following refractive index matching. We found that for two different species of echinoderms, *S. purpuratus* and the sea star *Patiria miniata*, large (>1 cm^3^) juveniles could be rendered nearly transparent using See-Star (Fig. 1D, E).

### Comparison of See-Star fluorescence microscopy performance with other clearing methods

To examine the utility of See-Star for fluorescence microscopy, we compared its performance to previously published clearing techniques. We prepared juvenile *P. miniata* using the See-Star protocol across a range of acrylamide concentrations (4, 15, and 30%) to compare how acrylamide concentration affects imaging depth. For comparison, we also prepared samples with other clearing protocols, including CUBIC and EZ-Clear, and with conventional preparations and mounting media, including 70% glycerol and 80% fructose (Fig. 2A). We stained all samples with DAPI, a highly permeable nucleic acid dye, to assess the impact of optical clarity on imaging depth. We hypothesized that differences in imaging quality would stem from variations in refractive index matching and/or depigmentation, influencing light penetration and scattering.

**Figure 2:**
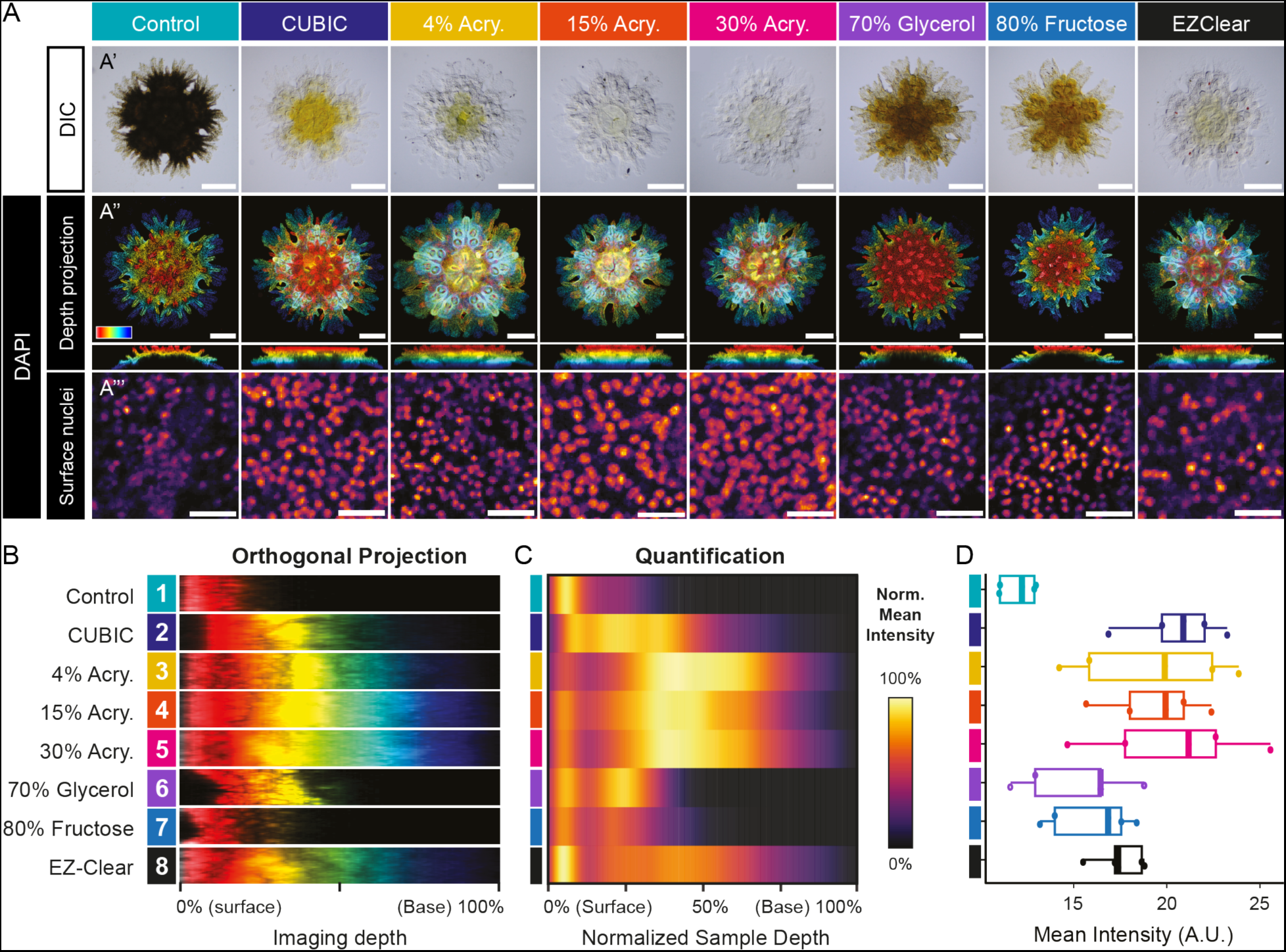
Comparison of See-Star fluorescence microscopy performance with other clearing methods A, Representative images of *Patiria miniata* juveniles following different clearing techniques visualized in DIC (**A’**) or in confocal microscopy after DAPI nuclei staining (**A’’**, **A’’’**). For DAPI-stained samples, color-coded (surface: red, base: blue) depth projections are imaged from the aboral side of the juveniles, with corresponding orthogonal views shown below (**A’’**). To clearly visualize the entire imaging depth, individual contrast adjustments were done for each condition. Close-ups on surface nuclei in representative areas are shown with identical contrast parameters (**A’’’**). In **A’**, **A’’**, scale bar = 200 µm. In **A’’’**, scale bar = 20 µm. **B.** Representative orthogonal projections from the center of the samples shown in **A’’**. **C,** Quantification of average sample intensity over normalized sample depth. n = 5 per condition. **D,** Quantification of mean DAPI fluorescence intensity within nuclei of surface epidermis following different clearing procedures. n = 5 per condition.

We found that the See-Star protocol, particularly at 15% and 30% acrylamide concentrations, provided superior optical clarity compared to other methods, followed by 4% acrylamide See-Star and EZ-Clear; these conditions outperformed conventional methods, which were still heavily pigmented following clearing (Fig. 2A’). Imaging depth was greatest in samples prepared using See-Star, and did not appear to vary across acrylamide concentrations (Fig. 2A’’, B, C). Both See-Star and EZ-Clear enabled imaging across the full depth of samples, unlike other methods where imaging was confined to surface layers (Fig. 2A’’, B). We also measured the normalized brightness of the DAPI signal throughout the samples, and found that peak signal intensity with See-Star occurred at approximately 50% depth, whereas for other methods, the highest intensity was near the surface, and decreased greatly with depth (Fig. 2C). Additionally, to evaluate the effect of mounting media on absolute signal intensity, we observed the brightness of nuclei at the surface, and found that See-Star and CUBIC produced the brightest signal, with negligible difference across acrylamide concentrations (Fig. 2A’’’,D).

### See-Star is compatible with immunohistochemistry in diverse marine invertebrates

Next, we tested whether See-Star was compatible with two of the most commonly used techniques for visualizing molecules, IHC and ISH. For IHC, we used a cross-reactive commercial anti-acetylated α-tubulin antibody, which allows for visualization of neurons and cilia in a broad range of invertebrate species (Formery et al., 2022; Herranz et al., 2020; Martín-Durán & Hejnol, 2015; Rawlinson, 2010; Santagata et al., 2012; Schwaha et al., 2018; Zieger et al., 2018). We first assayed this antibody in cleared *P. miniata* post-metamorphic juveniles, defined by bearing only two pairs of secondary tube feet in each ray (2-TF stage) (Fig. 3A). Neurite tracts within the circumoral nerve ring, the radial nerve cords as well as cilia on the epidermis, in the digestive tract and in the water vascular system were labeled, consistent with previous description of acetylated α-tubulin immunoreactivity in *P. miniata* at similar developmental stages (Formery et al., 2021). However, *P. miniata* post-metamorphic juveniles are small (< 1 mm) and can be easily decalcified by a short EDTA treatment to enable detailed microscopy. To test how See-Star enables whole-mount IHC in more difficult samples, we compare acetylated α-tubulin immunoreactivity of post-metamorphic juvenile stages with larger *P. miniata* juveniles at the 7-TF stage (∼ 5 mm) and 22-TF stage (∼ 1cm), which are much more opaque and calcified and would typically be impractical for whole mount imaging (Fig. 3B, C). At 7-TF and 22-TF stages, features similar to post-metamorphic stages were labeled, including the circumoral nerve ring, the radial nerve cords, and cilia associated with the digestive tract. Additional structures were labeled in the nervous system at these later stages. At the 7-TF stage, marginal nerves located on either side of the ambulacral regions (Cuénot, 1948) became clearly visible (Fig. 3B). In addition, series of nerves that we refer to as interradial nerves branched off regularly from the marginal nerves at the level of each tube foot, and extended laterally in the interradial areas towards the edges of the oral surface (Fig. 3B). The most proximal interradial nerves coalesced with their counterparts from the neighboring rays along the midline of the interradial area. Despite the large size of the specimens used, high-resolution imaging of smaller anatomical structures was still possible. For instance, we looked at the detail of the acetylated α-tubulin labeling in individual tube feet, showing that the lateral nerves branching off the radial nerve cords were prolonged along the stem of tube feet up to the base of the papilla, where a neurite network lined the entire epidermis, with a condensation facing the medial part of the arm (Fig. 3D,E). These observations are consistent with previous reports of tube feet neuroanatomy in various asteroid species (Elia et al., 2009; Formery et al., 2021; Moore & Thorndyke, 1993). Additionally, acetylated α-tubulin immunoreactivity also showed the presence of cilia inside the coelomic lining of the tube feet (Fig. 3E), as observed previously (Formery et al., 2021).

**Figure 3:**
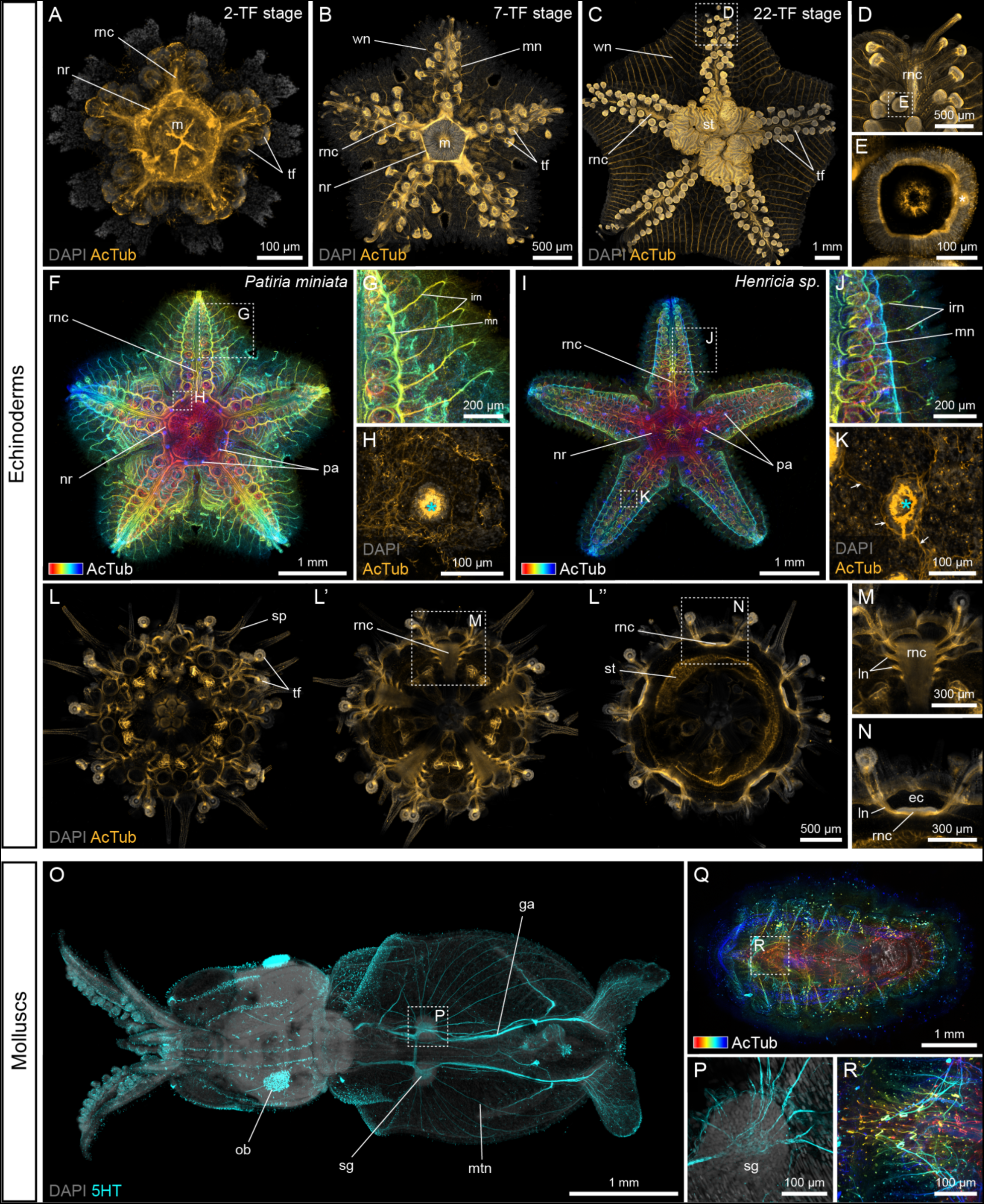
See-Star is compatible with immunohistochemistry. IHC in echinoderms (**A-N**) and molluscs (**O-R**) following See-Star clearing. **A-E,** Depth projections of acetylated α-tubulin IHC in 2-TF (**A**), 7-TF (**B**) and 22-TF (**C-E**) *Patiria miniata* juveniles. **D,** Magnification of the distal part of the arm outlined in **C**. **E,** Magnification of the tube foot papilla outlined in **D**. **F-K,** Comparison of α-tubulin IHC in a 11-TF *P. miniata* juvenile (**F-H**) and a 16-TF *Henricia sp*. juvenile (**I-K**). In **F**, **G**, **I**, **J**, acetylated α-tubulin staining is shown as a color-coded depth projection, so that the nerve ring on the oral side appears in blue while the papulae on the aboral side appear in blue. **G, J,** Magnification of the right side of the arms outlined in **F** and **I**, respectively. **H, K,** Magnification of an aboral sub-stack of the papulae outlined in **F** and **I**, respectively. In **H**, **K**, the cyan asterisks indicate the position of the papula. In **K**, the white arrows indicate condensations of basiepidermal neurites. **L-N,** Depth projections of acetylated α-tubulin IHC in a 11-TF *Strongylocentrotus purpuratus* juvenile. In **L-L’’,** distinct depth projections along the oral-aboral axis show the oral surface (**L**), the oral half (**L’**), and the equator (**L’’**) of the same juvenile. **M,** Magnification of the radial nerve cord outlined in **L’**. **N,** Magnification of the radial nerve cord outlined in **N’’**. **O, P,** Depth projection of serotonin IHC in *Doryteuthis opalescens* paralarvae. **P,** Close-up on the region of the stellate ganglion outlined in **O**. **Q, R,** Depth projections of acetylated α-tubulin IHC in a juvenile chiton (*Lepidozona sp.*). The staining is shown as a color-coded depth projection. **R,** Magnification of the sensory aesthetes outlined in **Q**. In **A-E, H, K-P**, nuclei are counter-stained with DAPI. ec: epineural canal; ga: giant axon; irn: interradial nerves; ln: lateral nerve; m: mouth; mn: marginal nerve; mtn: mantle nerve; nr: nerve ring; ob: olfactory bulb; pa: papula; rnc: radial nerve cord; sg: stellate ganglion; sp: spine; st: stomach; tf: tube foot.

We then validated the compatibility of See-Star with IHC in other echinoderm species by comparing acetylated α-tubulin immunoreactivity in cleared *P. miniata* with *Henricia sp*. juveniles (Fig.S1A), which belong to a distinct asteroid order (Fig. 3F-K). Both species exhibited similar neural organization, including the nerve ring and radial nerve cords, with marginal and interradial nerves displaying lateral branching at each tube foot. However, *Henricia sp.* had significantly shorter interradial nerves, aligning with its more compact interradial areas compared to *P. miniata* (Fig. 3G, J). A notable difference between the two species was the number of dermal papulae present on the aboral surface. In asteroids, papulae are outgrowths of the aboral coelom penetrating through the endoskeleton and protruding in the aboral epidermis (Fig.S2A), likely serving a respiratory function (Cobb, 1978). At the stages investigated, *P. miniata* juveniles only exhibited a pair of papulae at the proximal end of each arm, while *Henricia sp.* juveniles had a larger number of papulae spanning the entire surface of each arm (Fig. 3F, I; Fig.S2B, C). In both cases, the lining of the coelom forming the papulae was covered in strongly immunoreactive cilia. Closer examination of the papulae in *Henricia sp.* also revealed condensations of basiepidermal neurites around and between the base of the coelomic protrusion, whichwere not observed in *P. miniata* (Fig. 3H, K; Fig.S2D, E). In both cases, the samples were imaged from the oral side, while the papulae are located on the aboral surface, demonstrating the value of See-Star to allow high resolution morphological investigation across a thick layer of tissue.

To extend our study outside of asteroid species, we investigated the same antibody in 11-TF *S. purpuratus* juveniles. Unlike post-metamorphic echinoid juveniles that have a thin test and can be observed following short EDTA treatments and simple clearing methods (Formery et al., 2021; Thompson et al., 2021), older echinoid juveniles are extensively calcified and are completely opaque, offering a significant challenge for whole-mount imaging. Their fluid-filled cavity encased in a rigid test make them difficult samples for semi-thin sections as well, so that there are no straightforward methods for IHC on internal structures in this type of samples. Here, using See-Star we were able to observe for the first time the structure of the nervous system inside intact late echinoid juveniles (Fig. 3L-N), revealing the array of nerves innervating the spines (Fig. 3L), the circumoral nerve ring (Fig. 3L), the serial branching of the lateral nerves along the radial nerve cords (Fig. 3L’, M) and the epineural canal (Fig. 3L’’, N). These observations indicate that the general organization of the central and peripheral nervous systems at this stage are very similar to earlier description of post-metamorphic juveniles (Burke et al., 2006; Formery et al., 2021), although complexified by the higher number of appendages each provided with their own innervation. Altogether, our survey of acetylated α-tubulin immunoreactivity in asteroid and echinoid highlights the value of See-Star for comparing intact late developmental stages of large, opaque and highly calcified specimens that would otherwise only be accessible using destructive methods like dissection or semi-thin sections.

We next applied See-Star to two mollusc species to test the utility of the protocol outside of echinoderms in another phylum of heavily pigmented and calcified animals. First, to assess whether See-Star can adequately clear heavily pigmented samples, we attempted to visualize the nervous system of a cephalopod, the California market squid, *Doryteuthis opalescens*. Cephalopods have a variety of pigment-bearing cell types in the skin that enable the color-changing properties this group is known for (Cloney & Brocco, 1983). However, this abundance of pigment creates a challenge for observation of underlying structures by light microscopy. We found that *D. opalescens* paralarvae (∼5-6mm in length) processed with See-Star were nearly optically transparent, and a majority of the pigment had been removed (Fig. S1A). Using a commercial cross-reactive anti-serotonin (5HT) antibody, See-Star enabled visualization of the entire serotonergic nervous system of paralarvae (Fig. 3O), and also observation of fine structures, such as neural connections within the stellate ganglion (Fig. 3P). We also tested the utility of See-Star as a post-fixation on *D. opalescens* samples previously fixed in 4% paraformaldehdye and dehydrated in ethanol, and found that it performed equally well on freshly fixed samples and samples dehydrated for long term storage.

We reasoned that See-Star could also be applied to study structures that closely underly or pass through shells, such as the sensory aesthetes of chitons, which are neural sensory organs that reside in hollow channels within the valves (shell plates) (Boyle, 1976; Varney et al., 2024; Vendrasco et al., 2008). While heavily pigmented and calcified in life, chitons processed with See-Star were rendered nearly transparent (Fig. S1B). Anti-acetylated α-tubulin staining in juvenile chitons revealed for the first time the intact organization of the nervous system (Fig. 3Q), including neural processes in close proximity to the shell. We were able to observe the ultrastructure of aesthetes by confocal microscopy (Fig. 3R), which until now had only been observed by epoxy casting and electron microscopy.

### See-Star is compatible with colorimetric and chain reaction *in situ* hybridizations

ISH is another technique classically used to detect the localization of mRNAs, and which is amenable for non-model species, including in a wide range of marine invertebrates. To test whether See-Star is compatible with ISH, we used digoxygenin-labeled riboprobes to analyze the expression of the transcription factor *nkx2.1* in large (>1cm^2^) cleared *P. miniata* juveniles (Fig. 4A, B). We found *nkx2.1* to be expressed along the entire length of the radial nerve cords, which is consistent with expression patterns in earlier juvenile stages (Formery et al., 2023), but also confirmed the expression pattern predicted using RNA tomography at a similar stage (Formery et al., 2023). In addition, *nkx2.1* was also expressed in the digestive tract, which again was consistent with its expression in 2-TF stages (Formery et al., 2023). We then tested the compatibility of the See-Star clearing protocol with the Hybridization Chain Reaction (HCR) split-probe design, a new elaboration of fluorescent ISH (FISH) which uses short DNA probe pairs instead of full-length riboprobes, allowing cloning-free probe synthesis and versatile gene multiplexing (Choi et al., 2016, 2018). We used two different HCR FISH probe sets for *opn-4* and *myosin heavy chain* (*MHC*) in 8-TF *P. miniata* juveniles (Fig.4C-F). We found that *opn-4*, which code for a rhabdomeric photopigment (D’Aniello et al., 2015), was expressed at the tip on each arm in the optic cushion (Fig.4C, D). This is consistent with the presence in this organ of multiple photosensitive ocelli (Eakin & Brandenburger, 1979; Garm, 2017). In addition, we also found *opn-4* to be expressed in discrete cells located at the tip of each tube foot papilla (Fig.4E). Opsin expression in the tube feet has also been reported in echinoids (Ullrich-Lüter et al., 2011, 2013), reinforcing the idea that echinoderm tube feet are important sensory organs in addition to their locomotory role. In addition to *opn-4*, we looked at the expression of *MHC*, which is a terminal muscle differentiation marker expressed in striated muscles in all echinoderms (Andrikou et al., 2013; Perillo et al., 2023; Wessel et al., 1990). In *P. miniata* juveniles, we found this gene to be highly expressed in the water vascular system lining of the tube feet (Fig.4F), consistent with the presence of longitudinal muscle fibers in this tissue (Formery et al., 2021). *MHC* was also expressed in interradial muscles at the junction of the arms, and in the lining of the digestive tract (Fig.4F).

**Figure 4:**
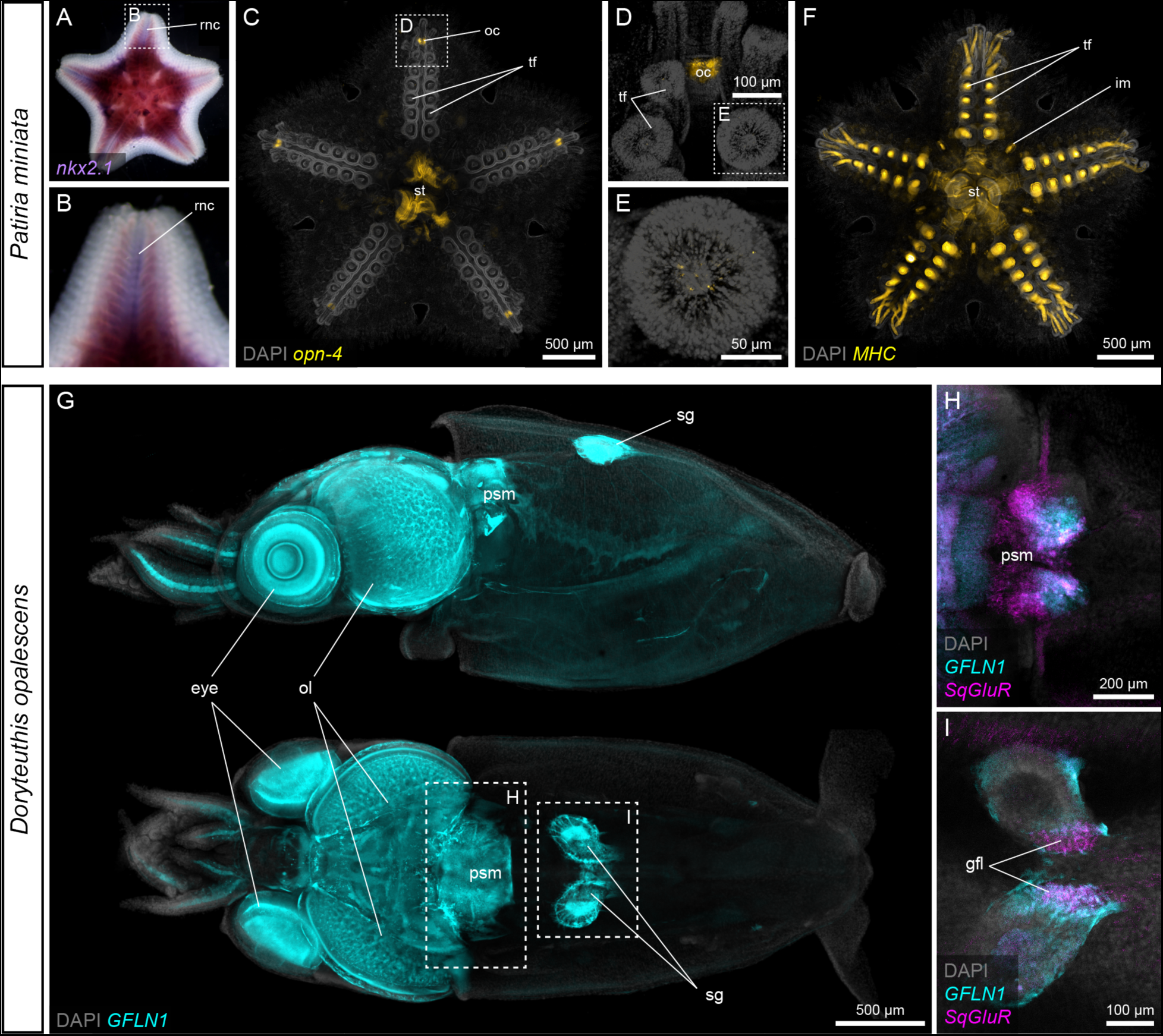
See-Star is compatible with *in situ* hybridization. ISH in *Patiria miniata* (**A-F**) and *Doryteuthis opalescens* (**G-I**) following See-Star clearing. **A, B,** Colorimetric ISH for *nkx2.1* in a large *P. miniata* juvenile. **B,** Magnification of the arm outlined in **A**. **C-E,** Depth projections of HCR FISH for *opn-4* in a 8-TF *P. miniata* juvenile. **D,** Magnification of the distal part of the arm outlined in **C**. **E,** Magnification of the tube foot papilla outlined in **D**. **F,** Depth projection of HCR FISH for *MHC* in a 8-TF *P. miniata* juvenile. Note that in **A-C, F,** background signal is present in the digestive tract. **G.** Depth projection of HCR FISH for *GFLN1* in a *D. opalescens* paralarva. Upper panel shows a lateral view, lower panel shows a dorsal view. **H, I,** Double HCR FISH for *GFLN1* and *Sq*G*luR* in a *D. opalescens* paralarva. **H,** Close-up on the brain region outlined in **G**. **I,** Close-up on the region of the stellate ganglia outlined in **G**. In **C-I**, nuclei are counter-stained with DAPI. gfl: giant fiber lobe; oc: optic cushion; ol: optic lobe; psm: posterior subesophageal mass; rnc: radial nerve cord; sg: stellate ganglion; st: stomach; tf: tube foot.

To validate that ISH is also compatible with the See-Star protocol outside of echinoderms, we designed HCR FISH probe sets in *D. opalescens* for two neural markers, the sodium channel *GFLN1* (Rosenthal & Gilly, 1993) and the glutamate receptor, *SqGluR* (Battaglia et al., 2003). We then investigated the expression of these genes in *D. opalescens* paralarvae (Fig.4G-I). The sequence for *GFLN1* was originally described from cDNA isolated from stellate ganglion extracts, and it was inferred to be expressed in the giant fin lobe of the stellate ganglion (Rosenthal & Gilly, 1993); however, its complete expression pattern was unknown. Here, using See-Star and HCR FISH, we find that *GFLN1* is highly expressed within the stellate ganglion, but also broadly in the brain and eyes (Fig. 4G). Our observations confirm that *GFLN1* is expressed in the cell bodies within the stellate ganglion that produce the giant fibers of the giant axon. *SqGluR* is specifically expressed in subregions of the brain, and in the giant fiber lobe of the stellate ganglion. *SqGluR* expression is enriched in the posterior of the brain, in a sub-region of the posterior subesophageal mass that is largely non-overlapping with expression of *GFLN1* (Fig. 4H). Within the stellate ganglion, *SqGluR* expression is restricted to the giant fiber lobe (Fig. 4I).

## Discussion

See-Star is a novel clearing method which effectively combines hydrogel crosslinking, chemical decalcification, and detergent-based delipidation into a single protocol to enable robust fixation and tissue clearing in large (>1 cm^3^), shelled and pigmented invertebrates, while preserving tissue integrity and morphology. The application of See-Star to large specimens, demonstrating deep tissue imaging and preservation of cellular structures, underscores its potential to push the boundaries of what can be studied in non-model organisms, but also in model organisms outside of the developmental stages which are typically surveyed. This is particularly relevant in the context of many marine invertebrates such as echinoderms or spiralians, in which embryos and larvae are often transparent, while later developmental stages corresponding to the formation of the adult body plan are much more challenging to image. See-Star is amenable for both heavily calcified and soft-bodied organisms, indicating that it is a versatile protocol that is useful for organisms of different body compositions and represents a significant advancement in the visualization of internal structures within calcified organisms, a long-standing obstacle in the field of evo-devo. Another key strength of See-Star, as demonstrated in our comparative microscopy studies, is its compatibility with both IHC and ISH, enabling the detailed visualization of protein and mRNA localization across entire organ systems. This compatibility allows for the exploration of gene expression patterns in stages of development previously inaccessible to whole mount imaging techniques, particularly in large juvenile and adult stages. In particular, See-Star’s compatibility with HCR FISH makes it highly suitable for comparative studies. Unlike IHC, which relies on the availability of target-specific primary antibodies, HCR FISH enables the exploration of expression patterns of any gene with readily available DNA probes which can be synthesized directly from cDNA sequences without molecular cloning, making it particularly attractive to use in emerging models. These properties make our See-Star protocol an invaluable tool to extend comparative studies in the field of evo-devo and provide new insights into the molecular mechanisms underlying animal body plans and organ systems.

There are several possible further optimizations that could be made to See-Star for tackling difficult tissues beyond what we tested. Consistent with other studies, we observed that depending on the samples, detergent-based delipidation did not always completely remove all light-absorbing pigments (Pende et al., 2020). This was for instance the case in the squid eye and chiton digestive tract (Fig.S1). While the small amount of pigment retained in some samples did not hinder our observations, in other heavily pigmented species it may be necessary to incorporate a bleaching step, such as H_2_O_2_ treatment (Konno & Okazaki, 2018; Pende et al., 2020). We also observed that background and autofluorescence can be an issue, especially in larger samples, so for some applications extended washing and addition of an autofluorescence quenching step may be necessary (Greenbaum et al., 2017). Lastly, we did not test whether endogenous GFP fluorescence is retained through the See-Star protocol. While this is not an issue in many non-model species where GFP transgenics are not available, it would limit the utility of the method for some model-system applications. However, based on the maintenance of GFP fluorescence through the CLARITY and Bone-CLARITY protocols that have similar chemistry to See-Star, it is likely that GFP would survive the See-Star process (Chung et al., 2013; Greenbaum et al., 2017).

In conclusion, See-Star is a powerful tool for comparative studies in a variety of fields, including developmental biology, neuroscience, and morphology, offering a new lens to examine the diversity of life. By enabling the detailed visualization of internal structures in non-model organisms, See-Star not only enhances our ability to conduct comparative studies but also opens up new possibilities for the investigation of molecular mechanisms and processes across a broader range of life history stages.

## Methods

### Animal collection

Adult specimens of *Patiria miniata* and egg cases of *Doryteuthis opalescens* were collected off the coast of Monterey Bay, California, US, and kept in circulating seawater tanks. For *P. miniata*, gravid adults were spawned in the laboratory by injecting 1 mL of 1 mM 1-methyladenine in each gonad. Following in vitro fertilization, *P. miniata* embryos and larvae were cultured at 14°C in UV-sterilized filtered seawater (FSW) at a density of about 1 per mL. The seawater was renewed every 2 or 3 days and the larvae were fed *ad libidum* with freshly grown *Rhodomonas lens* microalgae. After reaching metamorphosis, the juveniles were fed biofilm cultivated on circulating sea-water tables and supplemented by *R. lens* until they reached the appropriate size. For *D. opalescens*, paralarvae were hatched live from egg cases and maintained in flowing seawater. In addition, juvenile *Strongylocentrotus purpuratus*, *Henricia sp.*, and *Lepidozona sp.*, were collected directly at the appropriate size from the intertidal zone. All animals used in this study were collected with appropriate state permits.

### Fixation and clearing

A detailed See-Star protocol is provided in Supplementary Information 1. In short, animals were incubated in clean glass containers with FSW and without any food for at least two days and in some cases up to one week, in order to clear their gut from autofluorescent content. The samples were then anesthetized in a 1:1 mix of 7.5% MgCl and FSW on a flat petri dish before being fixed in 3.3X phosphate buffer saline (PBS) containing 4% paraformaldehyde and 30% acrylamide overnight at 4°C with gentle agitation. After the fixation, the samples were washed extensively in 1X PBS and incubated in gelation solution (1X PBS, 4% acrylamide, 0.25% VA-044) overnight at 4°C in septum-capped containers. Following the transfer in gelation solution and after the overnight incubation the containers were de-gassed using a syringe connected to a vacuum line. The gelation itself was performed subsequently by incubating the samples in gelation solution for 2 hours at 37°C. After gelation, the samples were carefully transferred to a decalcification solution (200 mM NaCl, 500 mM EDTA, 50 mM Tris-HCl at pH 8.5) and incubated at 37°C until any biomineralized structure had disappeared upon visualization under a stereoscope. This step typically took between 1 and 3 days to complete, and for highly calcified samples the incubation temperature was raised to 55°C. Once biomineralized structures were completely dissolved, the samples were transferred to a clearing solution (200 mM SDS, 200 mM NaCl, 50 mM Tris-HCl at pH 8.5) and incubated at 37°C until they became partially translucent. This step typically took between 1 and 3 days to complete, and for larger samples the incubation temperature was raised to 55°C. The samples were then washed extensively in 1X PBS and were ready for subsequent applications. Some of the samples used for IHC and ISH were initially fixed in 4% formaldehyde and dehydrated into 100% ethanol for long-term storage. These samples were rehydrated, post-fixed with acrylamide and cleared following the same procedure.

### Immunohistochemistry

After fixation and clearing, specimens were washed extensively with 1X PBS containing 0.5% Tween-20 (PBST) and then incubated for two hours in blocking solution (superblock (Thermofisher) supplemented with 0.05% triton X-100). The samples were then incubated for 24 to 48 hours at 4°C in blocking solution with the appropriate concentration of primary antibodies. The primary antibodies used in this study were a mouse anti-acetylated tubulin antibody (Sigma-Aldrich #T7451) diluted at 1:200 and a goat anti-serotonin antibody (Immunostar #20080) diluted 1:200. Following primary antibody incubation, the samples were washed extensively in PBS and then incubated overnight at 4°C in blocking solution with the appropriate concentration of secondary antibodies diluted at 1:500 and with DAPI (Invitrogen) diluted at 1:1000. The secondary antibodies used in this study were goat anti-mouse IgG and goat anti-rabbit IgG antibodies coupled to either Alexa488, Alexa546 or Alexa647 fluorophores (ThermoFisher). Following secondary antibody incubation, the samples were washed extensively in PBST, and then transferred to a refractive index matching mounting solution (50% weight/volume sucrose, 25% weight/volume urea, 25% weight/volume quadrol) modified from the CUBIC clearing protocol (Susaki et al., 2015).

### HCR fluorescent *in situ* hybridization

For HCR FISH, *P. miniata* and *D. opalescens* orthologues of *opn-4*, *MHC*, *GFLN1*, and *SqGluR* were identified from published literature (Battaglia et al., 2003; D’Aniello et al., 2015; Perillo et al., 2023; Rosenthal & Gilly, 1993) and short antisense DNA probe sets were ordered from Molecular Instruments using the full length cDNA sequence as input.

HCR FISH was performed as follows. After clearing and extensive washes in PBST, samples were permeabilized in detergent solution (1.0% SDS, 0.5% Tween-20, 150 mM NaCl, 1 mM EDTA, 50 mM Tris-HCl at pH 7.5) for one hour. The samples were then washed in PBST, and then in 5X saline sodium citrate buffer containing 0.1% Tween-20 (SSCT), before being pre-hybridized in hybridization buffer (Molecular Instruments) for one hour at 37°C. The probes were then added to the hybridization buffer at a final concentration of 0.05 µM and the samples were let to hybridize overnight at 37°C under gentle agitation. Following hybridization, the samples were washed 4 times 30 minutes in probe wash buffer (Molecular instruments) at 37°C and then in 5X SSCT at room temperature. They were then pre-amplified in amplification buffer (Molecular Instruments) for 30 minutes. Meanwhile, H1 and H2 components of the HCR amplifiers coupled to Alexa647 fluorophores (Molecular Instruments) were incubated separately at 95°C for 90 seconds, cooled down to room temperature in the dark and then pooled together before being added to the amplification buffer at a final concentration of 60 nM. The amplification was then performed overnight at room temperature. The samples were subsequently washed 4 times 30 minutes in 5X SSCT and incubated overnight in PBST containing DAPI (Invitrogen) diluted at 1:1000. Finally, the samples were washed in PBST and transferred to mounting solution before imaging.

### Colorimetric *in situ* hybridization

For colorimetric ISH, the *P. miniata* orthologue of *nkx2.1* was identified from published literature (Yankura et al., 2010), amplified from mixed stage *P. miniata* cDNA and cloned into pGEM-T Easy (Promega). Inserts were PCR amplified using M13 primers and template plasmid was digested with DpnI before PCR purification (Qiagen). Digoxygenin labeled antisense probes were synthesized using SP6 or T7 RNA polymerase (Promega), and purified using LiCl precipitation.

Colorimetric ISH was performed as follows. After clearing and extensive washes in PBST, samples were pre-hybridized in hybridization buffer (100µg.mL^-1^ heparin, 1 mg.mL^-1^ yeast RNA, 1X Denhardt’s, 5 mM EDTA, 0.1% CHAPS, 0.1% Tween-20, 50% formamide, 5X SSC) at 65°C for 4 hours. The samples were then incubated in hybridization buffer with 1 ng.µL^-1^ of probe overnight at 65°C. After hybridization, they were successively washed two times 1 hour in 2X SSCT at 65°C, three times 1 hour in 0.2X SSCT at 65°C, and three times 15 minutes in maleic acid buffer containing 0.5% Tween-20 (MABT) at room temperature. The samples were then blocked in blocking solution (maleic acid buffer containing 2% Boehringher-Mannheim blocking reagent) and incubated overnight at 4°C in blocking solution containing anti digoxigenin antibody (Roche) diluted at 1:2000. Following antibody incubation, the samples were washed extensively in MABT and then in alkaline phosphatase buffer (100 mM NaCl, 50 mM MgCl_2_, 0.1% Tween-20, 100 mM Tris-HCl at pH 9.5). The colorimetric reaction was performed by adding NBT/BCIP reagents (Roche) to the alkaline phosphatase buffer, and stopped at the appropriate time with MABT washes. Finally, samples were transferred to mounting solution before imaging.

### Image acquisition

Images of cleared samples before and after transfer to mounting solution and colorimetric ISH were taken with either a Zeiss discovery v12 stereoscope or a Zeiss Axioimager A2, in both cases using a Canon EOS T2i DSLR camera. Images of IHC and HCR FISH were taken using a Zeiss LSM700 confocal microscope. For confocal imaging, multichannel acquisitions were obtained by sequential imaging and large samples were acquired using the tile scan option.

### Image analysis and quantification

Confocal optical sections spanning regions of interest along the oral-aboral axis were compiled into maximum intensity z-projections using ImageJ v.1.52g. In some cases, a temporal color code was applied to visualize z depth. To quantify fragmentation, images were thresholded to produce a binary mask, and fragments were counted using the ‘Analyze particles’ tool in ImageJ. Fragmentation was calculated as the number of fragments divided by the total area of all fragments, then normalized on a zero to one scale. Transparency was calculated as the ratio of the difference in grayscale intensity values between black and white squares underneath samples divided by the same difference between squares outside of the sample. To quantify nuclear staining from fluorescent images, z-stacks of equivalent area were sub-sampled from the center of each image. Intensity of DAPI staining was then measured by segmenting nuclei using the ‘Analyze particles’ tool in ImageJ, and measuring average nuclear intensity per z slice. To quantify normalized fluorescence intensity across sample depth, intensity values and z-depth were normalized per sample, and then averaged within treatments. Absolute fluorescent intensity was measured from a surficial slice and averaged within treatments.

## Acknowledgements

We thank Lauren Lubeck, Paul Bump, and students in the Stanford BIOS236 course for help collecting animal specimens, and members of the Lowe laboratory for helpful discussions. This work was supported by an NSF grant (1656628) and Chan Zuckerberg BioHub funding to C.J.L.

## Author contributions

DNC and CJL conceptualized the work, and DNC developed the clearing protocol. DNC and LF conducted experiments and drafted the manuscript. CJL provided financial and intellectual support, and edited the text.

## Supplementary information

### See-star protocol

1. Let the specimens clear stomach contents for at least a couple of hours and ideally a couple days in clean filtered sea water (FSW), as microalgae are autofluorescent, and this signal can be maintained through the clearing process.
2. Anesthetize the specimens by incubating them in a 1:1 mix of FSW and 7.5% MgCl until they are completely relaxed.
3. **<u>Fixation step:</u>** **Note:** The fixation solution contains paraformaldehyde, which is a toxic chemical. The fixation step and subsequent washes should be performed in a fume hood with appropriate personal protection equipment and waste management.

1. Remove as much FSW as possible and incubate the samples in <u>fixation solution</u>.
2. For small samples (for instance embryos, larvae, or samples < 1 mm), fix for 2-4 hours at room temperature, with gentle rocking.
3. For medium samples (for instance squid paralarvae, small echinoderms juveniles, or samples < 5 mm^2^), fix overnight at 4°C, with gentle rocking.
4. For large samples (for instance large echinoderm juveniles, small chitons and bivalves, or samples < 10 mm^2^), fix for 24-48 hours at 4°C with gentle rocking. Not that larger samples are limited by mounting and imaging practicality for most microscopy set-ups. **Note:** The goal of this step is to allow for complete fixation and cross-linking between the acrylamide monomers and the proteins and nucleic acids within your sample. Thus, over fixation is not an issue; when in doubt about how long to fix, prefer a longer fixation time.
4. Rinse three times in 1X phosphate buffer saline (PBS) quickly, and then wash three times 20 minutes in 1X PBS
5. **<u>Gelation step:</u>**

1. Move the samples into a container that can be de-gassed (septum-capped vials, or septum-capped falcon tubes. Septa caps can be reused multiple times).
2. Remove as much PBS as possible and cover the samples using an excess of gelation solution, they should have at least ∼1 cm excess solution above them.
3. Acrylamide polymerization is inhibited by oxygen; degassing helps to mitigate this effect. De-gass the samples before the overnight incubation at 4°C:

1. Using a 22G syringe connected to an active vacuum line, pierce the septum cap to draw out the air in the container. Avoid sticking the syringe into the liquid. Gently
2. the vial on the edge of a bench helps release bubbles, and inserting and removing the syringe a few times can help maximize the amount of air removed.
3. The process is complete when there are no apparent bubbles in the sample vial.
4. If degassing is not feasible (e.g., if you don’t have septum-capped vials or no vacuum) you can still successfully gel your solution if you include excess gelation solution; the top several millimeters won’t gel, but everything below still should.
4. Incubate overnight at 4°C.
5. After overnight incubation, de-gass the sample vials again using the same procedure.
6. Incubate the samples at 37°C for 2 hours. Gelation is complete when the solution becomes viscous; a clear boundary should be barely visible between the polymerized solution around your sample and the layer of water or unpolymerized solution at the surface.
6. **<u>Decalcification step:</u>**

1. Rinse samples 3 times with PBS to remove gelation solution, then add <u>decalcification</u> <u>solution</u>.
2. Incubate the samples in decalcification solution at 37°C until they are completely decalcified. For large or highly calcified samples, this step can last several days. The decalcification solution should be changed daily. Increasing the temperature to 55°C speeds up the process, but can be damaging for fragile samples.
3. Check the samples regularly. Loss of refraction from the calcium crystals in shells or skeletal structures indicate that the process is complete.
7. **<u>Delipidation step:</u>**

1. Quickly wash the samples with <u>clearing solution</u>.
2. Incubate the samples in clearing solution at 37°C. As for decalcification, the speed of the process can be increased by incubating at 55°C. Gentle rocking is recommended if the samples are not fragile.
3. Check the samples regularly. Partial transparency and translucency in the tisses indicate that the process is complete. **Note:** the length, temperature and rocking of the clearing step have to be determined empirically by trial and error for different tissues and species. Prepare extra samples and plan to lose a few in the process; if you only have one precious sample, the safest condition is at 37°C with no shaking. Clearing is most rapid at 55°C with shaking, but at higher temperatures tissue warping and degradation are sometimes an issue.
8. Thoroughly wash the samples into 1X PBS.
9. **<u>Staining step:</u>** From that point, proceed to downstream applications (immunohistochemistry or *in situ* hybridization). **Note:** a few modifications to your downstream application protocols can be made, depending on the type of samples:

- For samples larger than 500 µm, at least double the length of your primary and secondary antibody incubations, and also increase the length and number of washes.
- Do not use phalloidin staining, actin has been denatured.
- Expect to troubleshoot antibodies: some antibodies don’t work as well in cleared tissues as they do with a standard PFA fixation (presumably due to denaturation of the tertiary structure of the epitope)
10. Following immunohistochemistry or in situ hybridization, move the samples in to CUBIC-mount (or another refractive index matching solution like glycerol).
11. Incubate overnight at 4°C in CUBIC-mount.
12. Wash the samples with a fresh volume of CUBIC-mount.
13. Mount your samples as desired for imaging. **Note:** coverslip-bottom dishes are very useful for larger samples

### Fixation solution

- 4% paraformaldehyde
- 30% Acrylamide
- 3.3X PBS (this corresponds to the salinity of seawater)

### Gelation solution

- 4% weight/volume acrylamide
- 0.25% weight/volume VA-044 (gelation agent)
- 1X PBS

Prepare using ice-cold reagents to avoid triggering the gelation process.

**Note:** excess Gelation Solution can be stored at 4°C for 1-2 days, or frozen at -20°C for several months. To check the stability of stored solutions, add 25 µL of Bis-Acrylamide to 975 µL of hydrogel solution (to a final concentration of 0.05%) and heat at 37°C for 2 hours; this solution should polymerize into a soft gel.

### Decalcification solution

- 200 mM NaCl
- 50 mM Tris
- 500 mM EDTA pH 8.5

Adjust to pH 8.5 with NaOH.

### Clearing solution

- 200 mM SDS
- 200 mM NaCl
- 50 mM Tris

Adjust to pH 8.5 with NaOH.

### Mounting solution

- 50% weight/volume sucrose
- 25% weight/volume urea
- 25% weight/volume quadrol

### Supplementary figures

**Figure S1:**
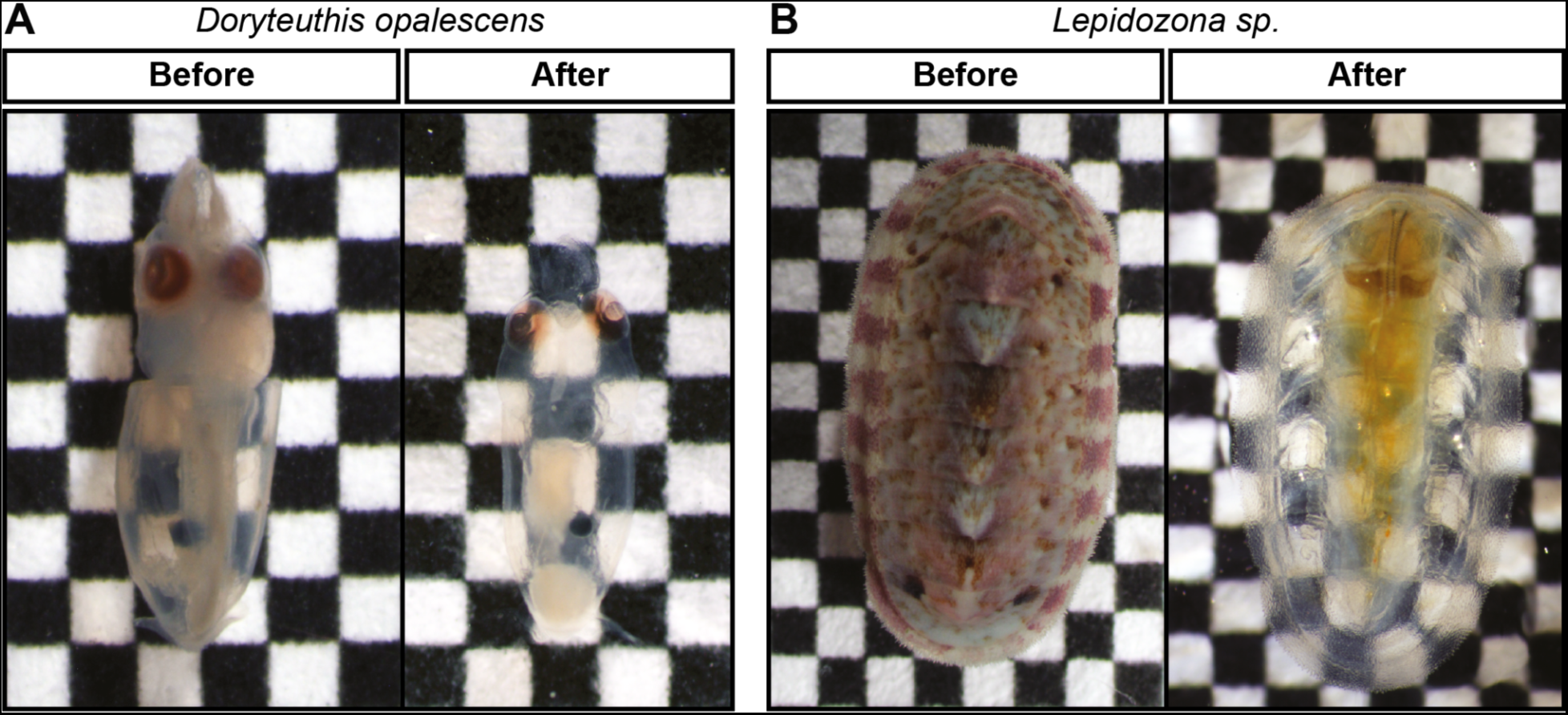
See-Star clearing of Doryteuthis opalescens and Lepidozona sp. Brightfield images of representative cleared *D. opalescens* paralarvae (**A**) and *Lepidozona sp.* (**B**) imaged before clearing (following fixation) and after clearing (before imaging). Squares = 600 µm.

**Figure S2:**
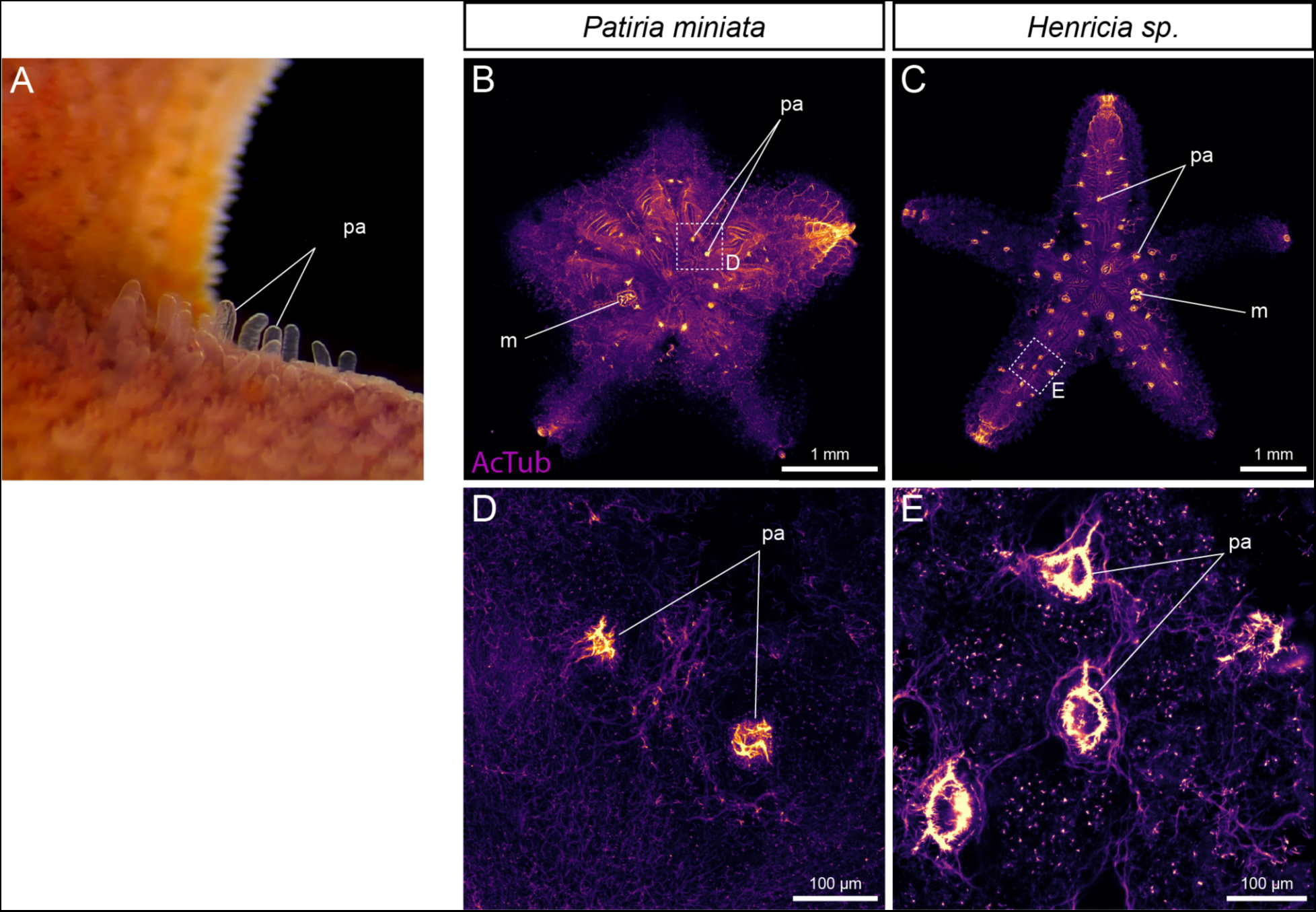
Papulae in *Patiria miniata* and *Henricia sp*. **A**, Brightfield image of the aboral surface of an arm in a live *P. miniata* juvenile, showing the protruding papulae. **B-E,** Acetylated α-tubulin IHC showing the aboral surface of a 11-TF *P. miniata* juvenile (**B, D**) and a 16-TF *Henricia sp*. juvenile (**C, E**). **D, E,** Magnifications of the regions outlined in **B** and **C**, respectively, showing details of the papulae. m: madreporite; pa: papula.

